# Impact of freeze-thaw-induced pit aspiration on stem water transport in a subalpine conifer (*Abies veitchii*)

**DOI:** 10.1101/2022.04.27.489725

**Authors:** Haruhiko Taneda, Mayumi Y. Ogasa, Kenichi Yazaki, Sachiko Funayama-Noguchi, Yoshiyuki Miyazawa, Stefan Mayr, Emiko Maruta

**Author notes:** **Corresponding Author:** H. Taneda, 7-3-1, Hongo, Bunkyo, Tokyo, Japan 113-0033, Japan.

## Abstract

During winter, subalpine conifers experience frequent freeze-thaw cycles in stem xylem, which may cause embolism and pit aspiration due to increased water volume during the sap to ice transition. This study examined the occurrence and ecological impacts of a combination of freeze-thaw-induced pit aspiration and embolism. In subalpine *Abies veitchii* trees, the fraction of closed pits and embolized tracheids as well as conductivity losses were measured to examine pit aspiration and its effects, triggered by natural and artificial stem freezing. When trees incurred mild drought stress in February and early March, 70% to 80% of stem conductivity was lost. Cryo-scanning electron microscopy indicated <20% embolized tracheids but ∼90% closed pits. Severe drought stress in late March caused 96 ± 1.2% (mean ± SE) loss of stem conductivity, while the fraction of embolized tracheids increased to 64 ± 6.6%, and aspirated pit fraction decreased to 23 ± 5.6%. Experimental freeze-thaw cycles also induced from 7.1 ± 0.89% to 49 ± 10% pit aspiration, and the fraction of closed pits was positively correlated to the percent loss of stem hydraulic conductivity. Results indicated that freezing-induced pit aspiration is an important factor for stem xylem dysfunction under mild drought. Upon severe drought in winter, stem water transport is predominantly inhibited by xylem embolism.

## Introduction

In evergreen conifers growing at high elevational habitats, winter cold leads to plant water deficiency due to the cease of water supply to leaves from frozen soil (i.e., frost drought; Mayr et al., 2012). In particular, trees at wind-swept habitats, such as at treelines, incur severe drought stress and, in consequence, lose stem hydraulic conductivity from early winter, often leading to the dieback of leaves and branches (Hadley & Smith, 1986; Maruta et al., 2020; Mayr et al., 2006; Sparks & Black, 2000). Xylem embolism, i.e., gas-filled tracheids inhibiting xylem water transport (Fig. 1a; Tyree & Zimmermann, 2002), is known to be one of the critical causes of xylem dysfunction in cold and harsh environments (Mayr et al., 2006; Ogasa et al., 2019; Maruta et al., 2020). The negative pressure, generated in dehydrated tissues and pulling the xylem sap as the driving force of plant water movement (Tyree & Zimmermann, 2002), can induce embolism, when gas bubbles enter a water-filled tracheid. Due to the strong negative xylem pressure in a drought-stressed plant, gases in an embolized tracheid can spread into neighboring water-filled tracheids via adjacent pit-membranes, thereby blocking xylem sections (Bouche et al., 2014; Sperry et al., 1996; Sperry & Tyree, 1990).

**Figure 1.**
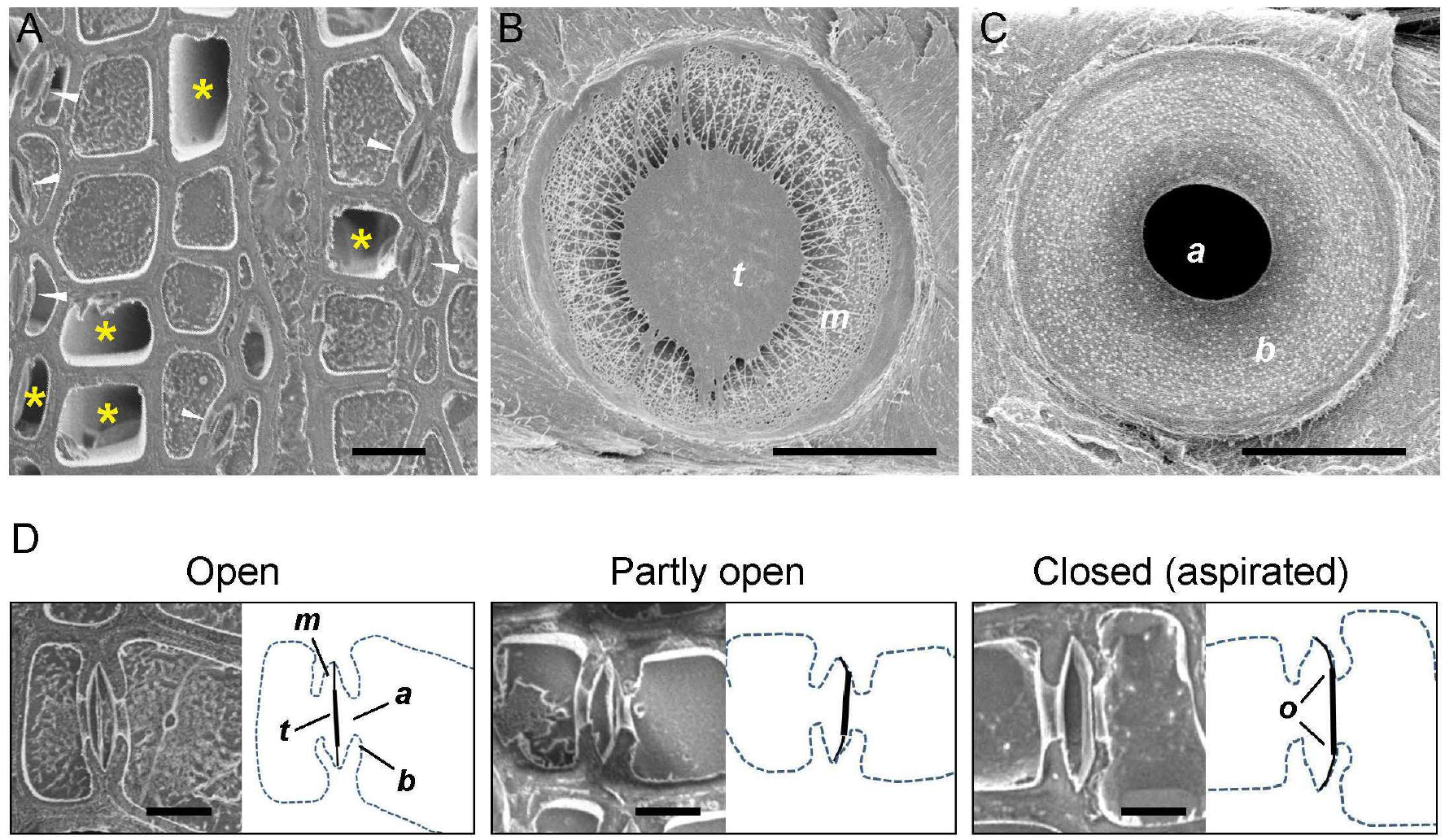
Pit-membrane of *A. veitchii*. (A) Water- and gas-filled tracheids in the xylem. (B) Torus-margo structure of a pit-membrane. (C) Pit aperture and warty layer on the pit border. (D) Three positions of a pit-membrane in a pit: open, partly open, and closed. (A) Cryo-SEM observations on the cross-section of the stem xylem in *A. veitchii*. Asterisks represent gas-filled (embolized) tracheids, and arrowheads represent exemplary pits. A linear structure in a pit is a pit-membrane. Scale bar, 10 μm. (B and C) After freeze-drying for 3 days, the 1-year-old stem xylem was split and coated on the newly appeared surface with an osmium coater and observed with a field-emission SEM (Sano, 2005). (D) The right illustration of each panel is a tracing of the left images. Solid lines are pit membranes, whereas a thick line in the central part represents a torus. Dashed lines are the outlines of tracheid walls. Letters in panels are as follows: a, pit aperture; b, pit border; m, margo; o, the overlap between the pit border and the torus; t, torus. Scale bar, 5 μm (B–D).

Additionally, freezing of the xylem sap causes the formation of gas bubbles entrapped in ice (Sperry & Sullivan, 1992; Utsumi et al., 1998; Sevanto et al., 2012). When ice thaws, small gas bubbles dissolve back to the sap because the surface tension between the water and the bubble forces the gas into solution. Large gas bubbles can expand and fill the tracheid lumen by the xylem negative pressure. As the gas volume in the sap is related positively to conduit volume, freeze-thaw cycles can cause severe xylem embolism in addition to drought stress-induced embolism in stems bearing wide and long conduits (Davis et al., 1999; Pittermann & Sperry, 2006; Willson & Jackson, 2006). Subalpine conifers that contain narrow-diameter tracheids (10–20 μm; Pittermann & Sperry, 2006; Mayr et al., 2006,) are potentially resistant to freeze-thaw-induced embolism (Feild & Brodribb, 2001; Pittermann & Sperry, 2003; Sperry & Sullivan, 1992). However, frequent freeze-thaw cycles were demonstrated to increase stem conductivity losses even in a comparatively mild drought condition during winter (Mayr et al., 2003, 2007; Sparks et al., 2001; Sparks & Black, 2000).

Recently, several studies have suggested that pit closures due to pit aspiration were associated with winter xylem dysfunction related to repeated freeze-thaw events (Mayr et al., 2014; Mayr et al., 2019; Maruta et al., 2022). Pit aspiration in conifer tracheids is commonly found when embolism occurs (Tyree & Zimmermann, 2002). Conifers possess pit membranes consisting of a thick-walled “torus” in the central part and a fibrous “margo” in the periphery (Fig. 1B and C; Hacke et al., 2004; Jacobsen, 2021). When cavitation occurs in a tracheid, the large pressure difference forms between neighboring gas- and water-filled tracheids, which allows the torus to move towards the cell wall of the pit border due to the flexible structures of the margo and to close the pit aperture (i.e., pit aspiration; Fig. 1D, bottom right; Sperry & Tyree, 1990; Hacke et al., 2004; Zenlinka et al., 2015). The pit closure by the aspiration prevents the further spread of gases to the adjacent tracheids. Theoretically, pit aspiration may also occur on freezing of the xylem sap in stems (Hammel, 1967). The dynamics and spatial patterns of freezing in stem tissues are complex: ice nucleates at some points and then propagates into other parts of the xylem in both axial and radial directions (Neuner et al., 2010; Charrier et al., 2017). When ice spreads through the xylem, the increment of ice volume by ∼8% may directly move pit membranes. Alternatively, its expansion may push the unfrozen water and generate a pressure difference and water flow along unfrozen water-filled tracheids across pits, resulting in pit aspiration. If pit membranes are retained at the aspirated position (Fig. 1D), the closing of the pit aperture may reduce hydraulic conductivity in the absence of embolism.

Pit aspiration of tracheids was found in needles and/or stem xylem of *Picea abies* (Mayr et al., 2014), *Pinus mugo* (Mayr et al., 2019), and *Abies mariesii* (Maruta et al., 2022) during winter when stems experience subfreezing temperature. In *P. abies*, the fraction of aspirated pits was related positively to loss of stem conductivity and the number of freeze-thaw cycles (Mayr et al., 2014), whereas the related studies also demonstrated the occurrence of xylem embolism via the acoustic emission (Mayr et al., 2007; Mayr & Sperry, 2010) and cryo-scanning electron microscopy (cryo-SEM) method (Mayr et al., 2007). The observed pit aspiration might be caused by freezing of xylem sap as well as by occurrence of embolism. In contrast, Maruta et al. (2022) detected the pit aspiration without embolism by cryo-SEM and suggested that winter xylem dysfunction was elucidated predominantly by sap-freezing induced pit aspiration rather than embolism in *A. mariesii*. Winter xylem dysfunction may thus be caused by embolism or pit aspiration, or a combination of both.

This study examined the contributions of pit aspiration and xylem embolism induced by sap freezing to stem water transport under both field and experimental conditions on subalpine conifer *Abies veitchii* Lindl., a dominant species in subalpine forests of the main island of Japan (Tsuyama et al., 2015). For field measurements, seasonal loss and regaining of stem hydraulic conductivity of 1-year-old internodes were monitored from winter to summer. On the stem segments, the distribution of air- and water-filled tracheids and the fraction of closed pits were observed using cryo-SEM (Fig. 1). The occurrence of pit aspiration due to sap freezing rather than xylem embolism should be detected as closed pits between water-filled tracheids (Maruta et al., 2022). In the laboratory, to demonstrate that sap freezing causes the occurrence and retaining of pit aspiration, 1-year-old internodes of *A. veitchii* were subjected to mild drought, followed by experimental freeze-thaw cycles, enabling the monitoring of pit aspiration during consecutive frost cycles. Further, an injection of water into a stem segment under increasing and decreasing pressure was conducted. If the closed position of the pit-membrane is retained in water-filled tracheids, stem conductivity reduced under higher applied pressure is maintained after the applied pressure is relaxed. This experiment should also enable insights into the mechanical flexibility of the margo and demonstrate pressure thresholds for the movement of the torus and pit aspiration. Based on field and experimental data, this study assessed the occurrence of freezing-induced pit aspiration and the resulting hydraulic blockage *in natura* and analyzed its relevance for hydraulic limitations in combination with xylem embolism and for hydraulic recovery in *A. veitchii*.

## Results

### Xylem dysfunction and pit aspiration in the field

Wind-exposed branches of *A. veitchii* trees incurred severe drought stress (Fig. 2B and C). Shoot water potential dropped from −1.8 ± 0.22 MPa in early February to −2.6 ± 0.12 MPa in late March (Fig. 3A). After the first rain event in April, shoot water potential recovered significantly to −0.54 ± 0.06 MPa and reached ∼0 MPa in late July. Percent loss of stem hydraulic conductivity (PLC) increased from 73% ± 7.8% to 96% ± 1.2% between early February and late March (Fig. 3B), gradually decreased to 55% ± 4.5% from April to June, and rapidly dropped to 17% ± 3.1% in late July. PLC between May and July was significantly lower than those of late March. As the new xylem growth started in late June (Fig. 4G), xylem recovery was thus not related to xylogenesis.

**Figure 2.**
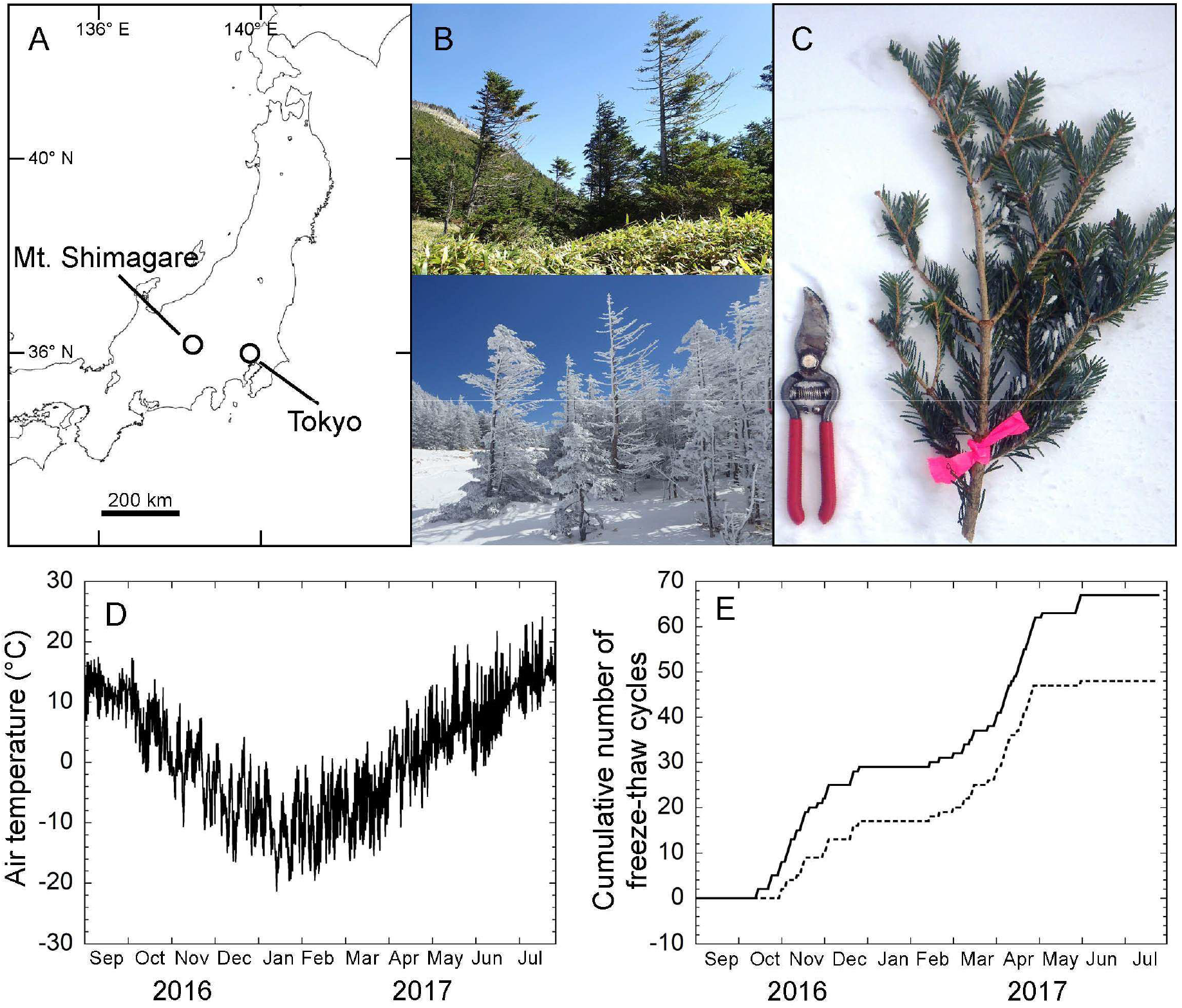
Study site and plant materials. (A) Sampling site located at Amaike Pass. (B) Wind-exposed trees that grew on the study site in September 2016 (top) and February 2017 (bottom).(C) Sampled branch with mechanical wounds. (D) Seasonal changes in air temperature from September 2016 to July 2017. (E) Cumulative number of freeze-thaw cycles. Solid and dotted lines represent the cumulative number of freeze-thaw cycles for xylem sap freezing at −1°C and −2°C, respectively.

**Figure 3.**
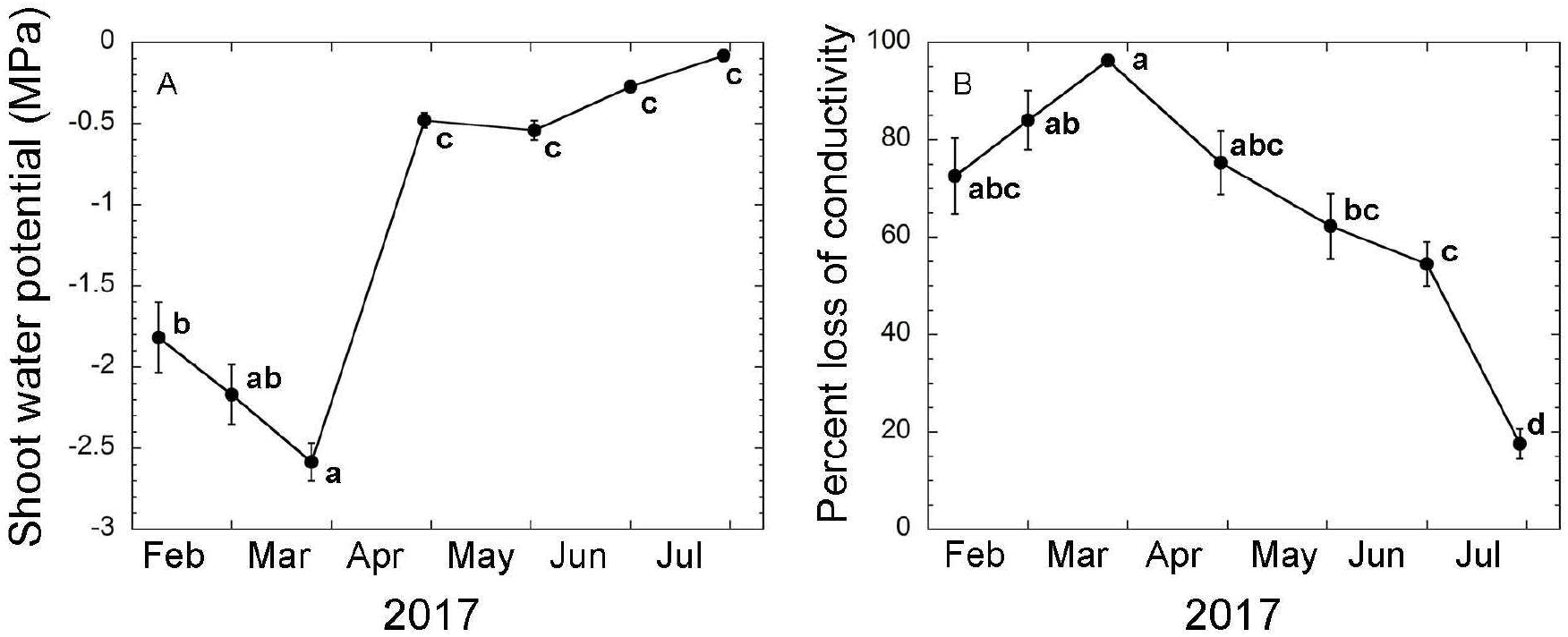
Water status and stem water transport of *A. veitchii* trees from February to July 2017. (A)Water potential of a current-year shoot. (B) PLC in the 1-year-old internode measured by a hydraulic method. The radial growth of the stem xylem started in late June. Error bars represent ±1 SE (*n* = 5–6). Different letters indicate significant differences at *P* < 0.05 (ANOVA followed by Tukey’s correction for multiple pairwise comparisons).

**Figure 4.**
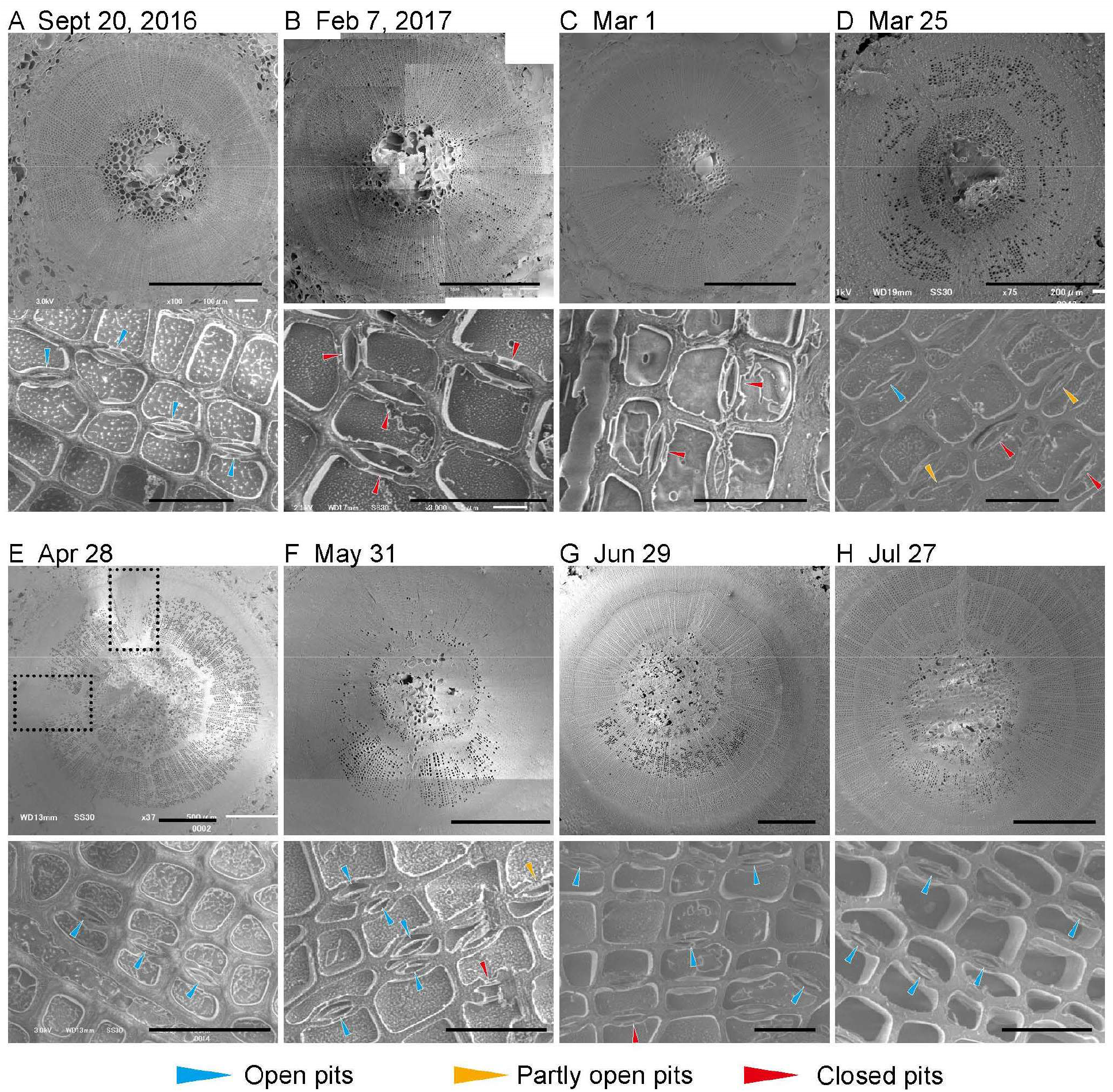
Seasonal changes in the distribution of air-filled conduits and pit-membrane position between September 2016 and July 2017 by cryo-SEM observations. Images represent (A) September 20, 2016, (B) February 7, 2017, (C) March 1, 2017, (D) March 25, 2017, (E) April 28, 2017, (F) May 31, 2017, (G) June 29, 2017, and (H) July 27, 2017. The normal and reaction woods were exhibited at the top and bottom parts of the image of the stem cross-section, respectively. Blue, yellow, and red arrowheads are open, partly open, and closed positions of the pit membrane, respectively. Square regions outlined by dotted lines in (E) represent leaf traces. Scale bar, 500 μm (top) and 20 μm (bottom).

Cryo-SEM observations enabled insights into the distribution of water- and air-filled tracheids within stem xylem: Stems sampled in September 2016 contained no embolized tracheids (Fig. 4A, top). Notably, in February and early March 2017, solitary embolized tracheids were sparsely distributed in the 1-year-old xylem (Fig. 4B and C, top). The embolized tracheids were found mainly in reaction wood but occupied on average <20% of the entire xylem (Fig. 5A). In late March, numerous tracheids were embolized (Fig. 4D, top). The percentage of embolized tracheids significantly increased up to 64% ± 6.6% of the entire xylem, whereas 85% ± 4.2% of tracheids were embolized in reaction wood (Fig. 5A). Between April and July, the percentage of embolized tracheids gradually decreased from 33% ± 5.8% to 5.1% ± 2.6% (Fig. 5A). Embolized tracheids in reaction wood were found at an overall higher fraction than in normal wood in this period, but the decrease showed similar dynamics (Fig. 5A; Table S1).

**Figure 5.**
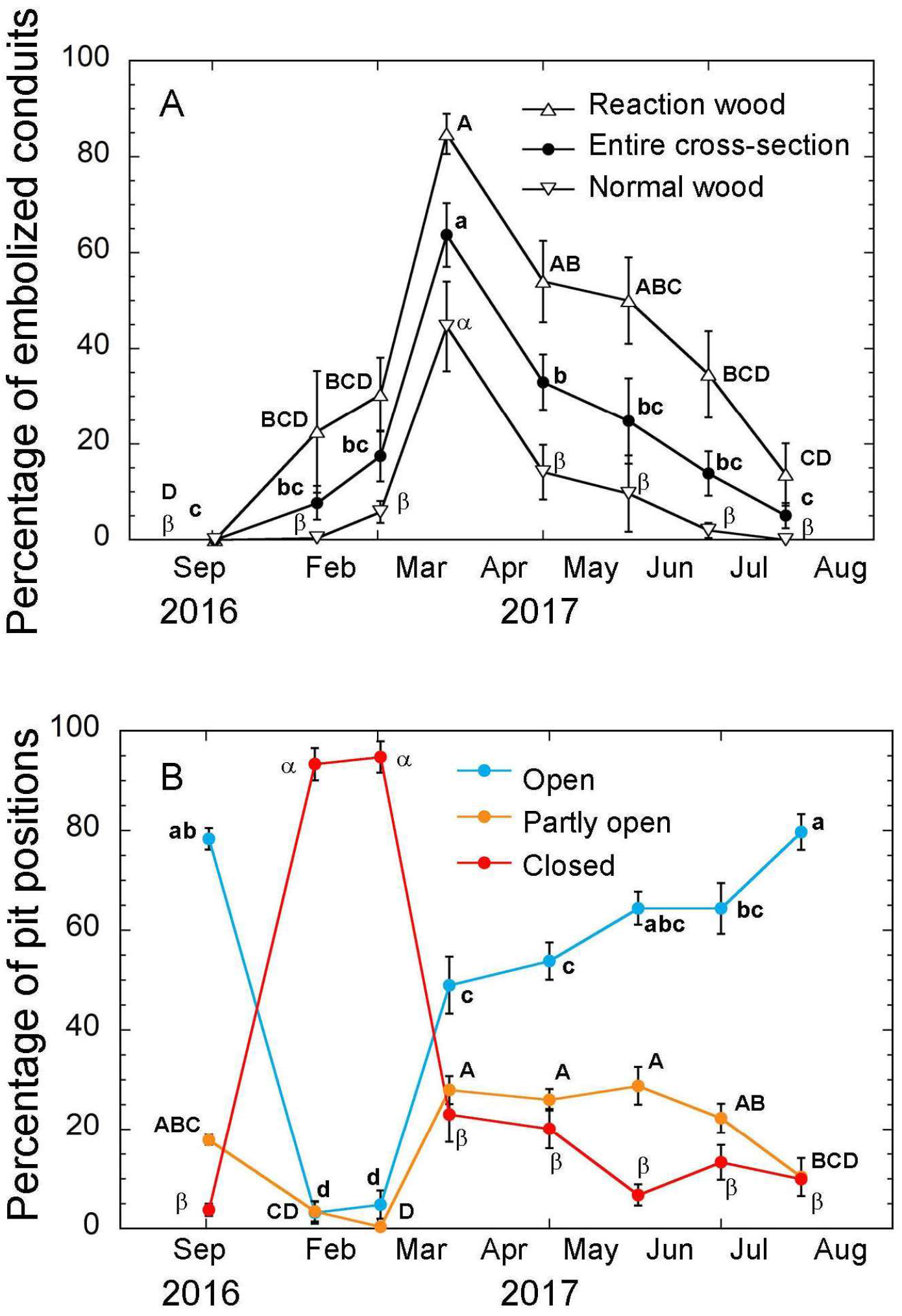
Seasonal changes in fractions of embolized tracheids and pit-membrane positions based on cryo-SEM observations in wind-exposed stems. (A) A fraction of the number of embolized tracheids. Black circles represent the average value among the four parts of the xylem. The top and bottom directed triangles represent the fractions in reaction and normal wood. (B) Fractions of the pit-membrane position. Blue, yellow, and red circles represent open, partly open, and closed pits. Error bars represent ±1 SE (*n* = 4–7). Different letters indicate significant differences at *P* < 0.05 (ANOVA followed by Tukey’s correction for multiple pairwise comparisons).

Cryo-SEM observations also allowed for detailed analyses of pit-membrane positions (Fig. 5B; Table S2). In September 2016, most tracheid pit membranes were located in a central position (i.e., open pits; Fig. 4A, bottom; Fig. 5B). In February and early March 2017, >90% of the pit membranes were located in an aspirated position, and adjacent tracheids were always filled with water (Figs. 4B and C and 5B, bottom). In late March, the fraction of the closed pits rapidly decreased to ∼20%, whereas about half of the pits were in an open position (Fig. 5B). The fraction of open pits gradually increased from April to July and reached 80% ± 3.6%.

### Effect of freeze-thaw events on hydraulic dysfunction and pit aspiration

Vulnerability curves obtained by dye-perfusion method indicated that *P*_50_area_, which is the xylem water potential at which 50% of conductive area is lost, increased with the number of freeze-thaw cycles. *P*_50_area_ on drought treatment was −2.8 ± 0.22 MPa but −2.0 ± 0.17 and −1.3 ± 0.053 MPa after drought combined with 10 and 15 frost cycles, respectively (Fig. 6). Native PLC and water potential measured in February and March corresponded to vulnerability curves of combined drought and freezing (Fig. 6, large open circles).

**Figure 6.**
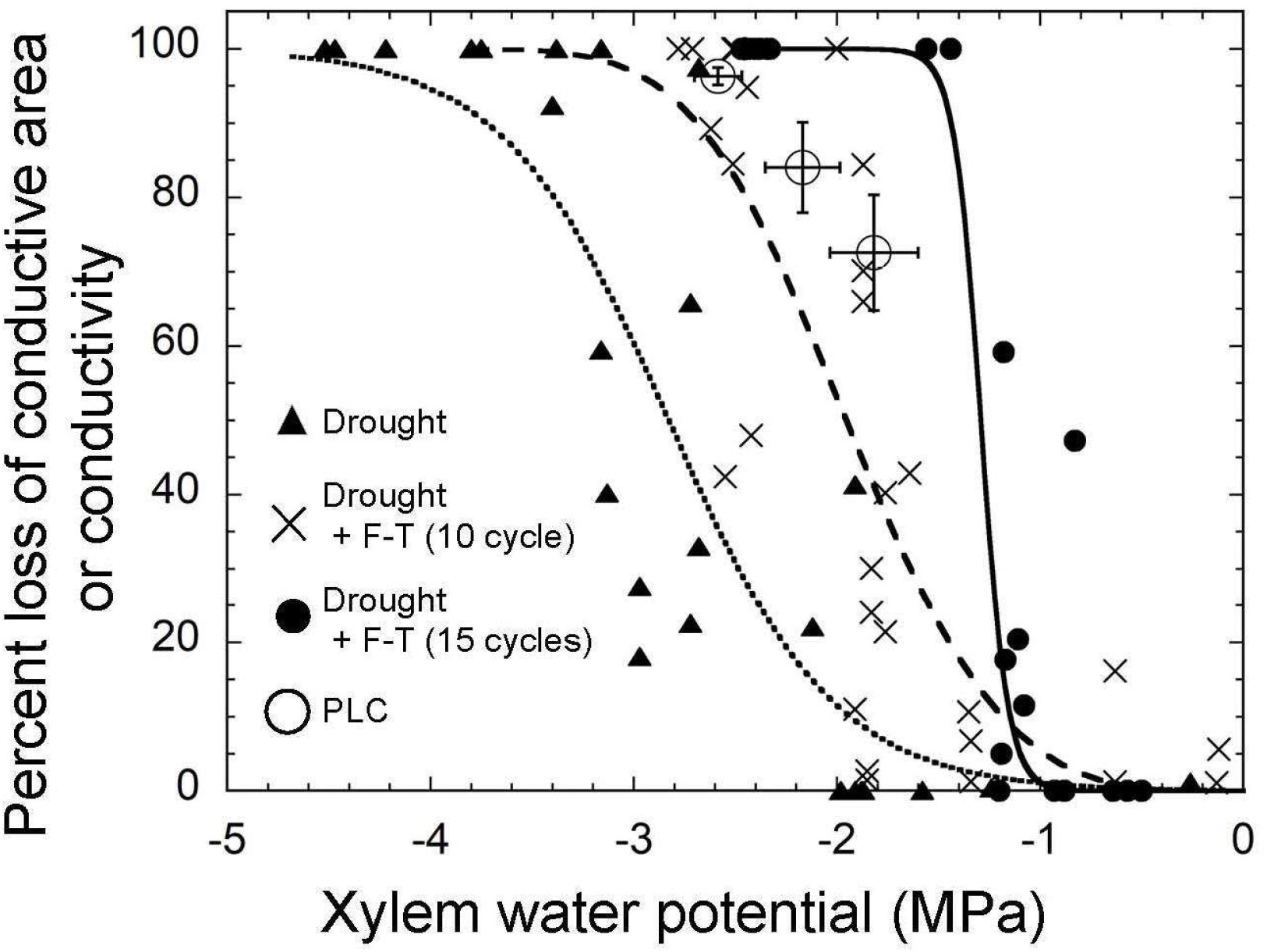
Comparison of vulnerability to drought- and freeze-thaw-induced embolisms in 1-year-old internodes of *A. veitchii* stems. The relationship between PLC_area_ (determined by the dye perfusion method) and xylem water potential was measured in stems subjected to drought stress followed by 15 freeze-thaw cycles (filled circles and a solid curve), drought stress followed by 10 freeze-thaw cycles (crosses and a dashed curve), and drought stress (filled triangles and a dotted curve). The latter two data were derived from Ogasa et al. (2019). Large open circles are the native PLC (determined by the hydraulic method) of stems sampled from field-growing trees in February and March 2017. Error bars represent ±1 SE (*n* = 6–9).

Before experimental freeze-thaw cycles, 79% ± 1.9% of pit membranes were in an open position and only 7.1% ± 0.89% of pits were closed (Fig. 7A). After freeze-thaw cycles, 49% ± 10% of observed pits were closed, whereas the fraction of open pits decreased to 37% ± 6.8%, whereby the percentage of closed pits was positively correlated to PLC (Fig. 7B). Stem segments regained 75% ± 3.4% of open pits after submersion in water under a partial vacuum overnight (Fig. 7A).

**Figure 7.**
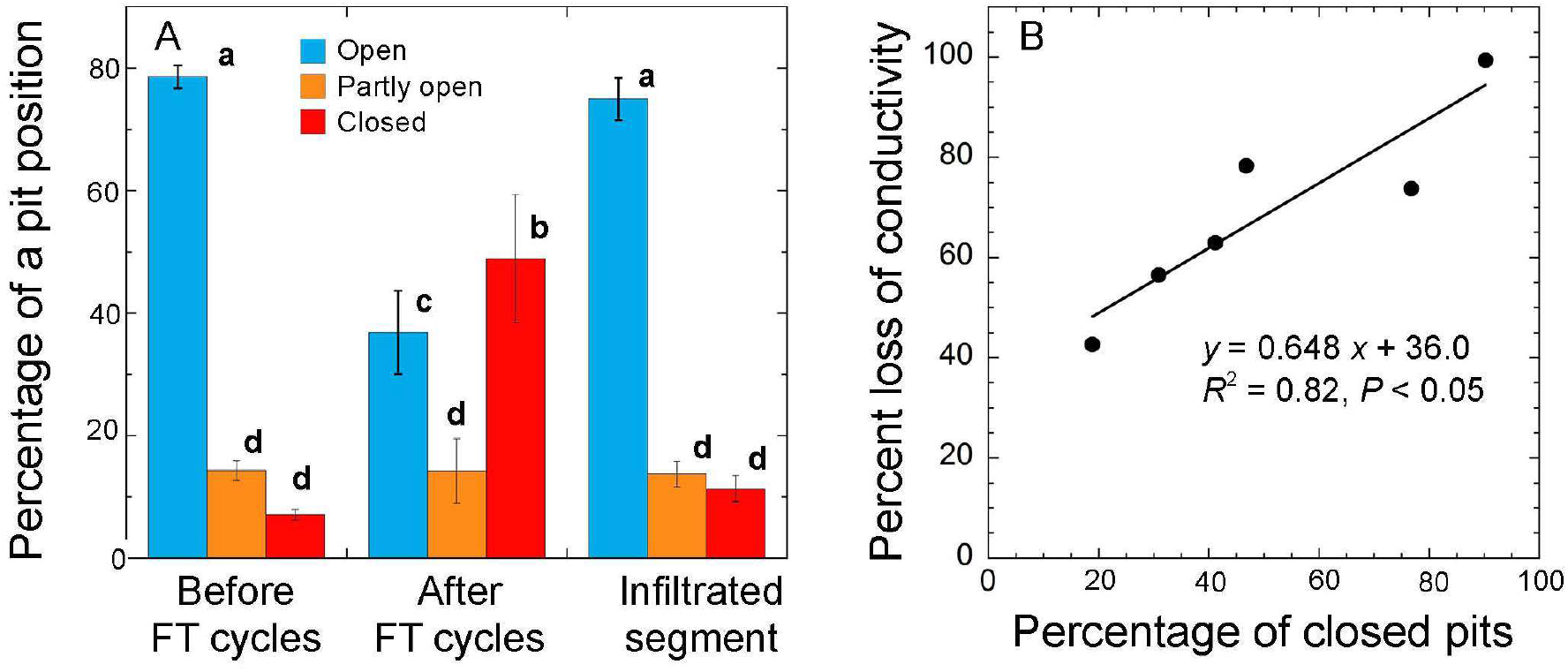
Effect of experimental freeze-thaw cycles of the xylem sap on the pit-membrane position of a tracheid in stems dried to −2.0 MPa. (A) Changes in the pit-membrane position before and after freeze-thaw treatment of the xylem sap. Right columns represent the pit-membrane positions of stem xylem soaked in a partial vacuum overnight after freeze-thaw treatment. Blue, yellow, and red columns are open, partly open, and closed pits (see Fig. 1D). (B)Relationship between PLC and the percentage of closed pits in stems with freeze-thaw treatment. Error bars in (A) represent ±1 SE (*n* = 6). Different letters indicate significant differences at *P* < 0.05 (ANOVA followed by Tukey’s correction for multiple pairwise comparisons). Data in (B) are regressed with a linear function.

### Pit aspiration induced by water injection

The water flow inducing pit aspiration was measured by changes in hydraulic conductivity, during gradually increasing pressure. Stem conductivity started to decrease when water flow through a stem segment was generated at the applied pressure of >1.22 ± 0.059 kPa/mm (at 95% of the maximum conductivity) and dropped to 50% of the maximum value at 3.65 ± 0.089 kPa/mm (Fig. 8). Stem conductivity remained low even when the applied pressure was relaxed. A control experiment with *Abies sachaliensis* stems confirmed that water flow at high-pressure (=250 kPa and 6.25 kPa/mm) caused ∼80% reduction in stem conductivity and a high fraction of closed pits (Fig. S2).

**Figure 8.**
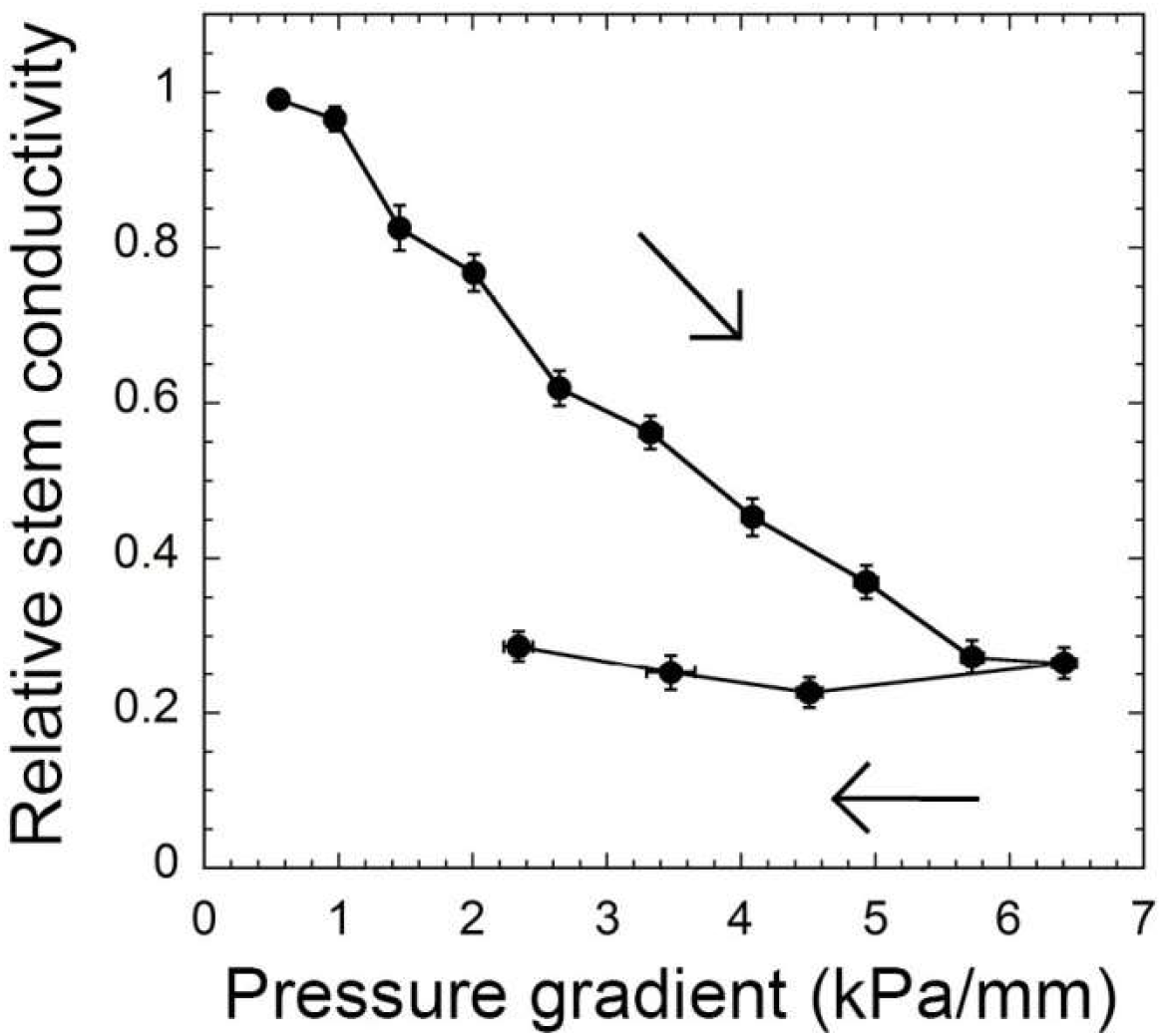
Reduction in stem conductivity of 1-year-old internodes of *A. veitchii* stem in water flow at a high-pressure gradient. The relationship between relative stem conductivity and a pressure gradient driving a flow was shown. Arrows represent an order to apply pressure to the stem segment. Error bars represent ±1 SE (*n* = 5–6).

## Discussion

Pit aspiration occurred both *in natura* and when branches were experimentally exposed to freeze-thaw cycles. At the treeline, pit aspiration upon sap freezing caused a substantial reduction in stem hydraulic conductivity, even in the absence of embolism and before severe drought stress-induced embolism in late winter. Pit aspiration may thus amplify xylem hydraulic dysfunction caused by embolism, and a combination of pit aspiration and embolism cause long-term hydraulic limitation in subalpine conifers (Mayr et al., 2006; Maruta et al., 2020). The reduced stem conductivity due to freezing-induced pit aspiration was also confirmed in artificial freeze-thaw treatments. Finally, water-injection experiment suggested that the aspirated pit membrane did not elastically return to a normal position and was retained for a while.

### Ecological impacts of pit aspiration induced by sap freezing (field observations)

In February and early March 2017, when trees were subjected to mild drought, pit aspiration was the major factor impairing water transport in *A. veitchii* stems. During this period, closed pits between water-filled and thus hydraulically functional tracheids were frequently found within the xylem (Figs. 3 and 5B), indicating that freezing events promoted pit closure. The aspirated position of pit membranes was obviously maintained after thawing of the xylem sap. Similarly, in *P. abies* and *A. mariesii* stems, increasing fractions of closed pits were detected during winter, whereas the xylem water potential was moderate (Maruta et al., 2022; Mayr et al., 2014). It is likely that freezing-induced pit aspiration substantially contributed to xylem dysfunction in the two subalpine conifers, although *P. abies* stems also showed some embolism (Mayr et al. 2007). During early winter when freeze-thaw cycles were frequently recorded (i.e., October to early January in Fig. 2D and E; also see Mayr et al., 2007; Maruta et al., 2020), freezing-induced pit aspiration may be a relevant factor limiting xylem hydraulic function and impairing stem water transport in plants under moderate drought stress. Interestingly, a high fraction of closed pits in early winter was also detected in *P. mugo* (Mayr et al., 2019) but, in contrast to *A. mariesii, A. veitchii* and *P. abies*, this did not correspond to high PLC. The discrepancy might be explained by a species-specific deaspiration of pits due to the weak positive pressure during conductivity measurements. We suggest that aspirated pits of *P. mugo* show a weaker adhesion than the two *Abies* species. and *P. abies*.

In February and early March 2017, solitary embolized tracheids were found only sparsely within the xylem; in late March, severe xylem embolism contributed strongly to the observed decrease in hydraulic conductivity (Figs. 3–5). Solitary embolized tracheids found in February and early March were likely derived from gas bubbles generated during freeze-thaw cycles (Sevanto et al., 2012). When the xylem water potential was as negative as *P*_50_area_ for drought-induced embolism in late March (Figs. 3A and 6), these solitary embolized tracheids may have been starting points for air seeding and the formation of larger embolized xylem areas as observed during late winter (Fig. 5A; see also Mayr et al., 2007; Mayr & Sperry, 2010). In late March, the fraction of embolized tracheids increased to ∼60% and PLC increased nearly to 100% (Fig. 3B). The embolized tracheids were more frequently found in earlywood (relative to latewood; Fig. 4D and E), which contributes more to conductivity (or loss of conductivity when embolized). Thus, the high PLC was a result of embolism combined with pit aspiration.

Furthermore, PLC decreased to <20% until July, similar to seasonal courses of winter embolism observed in European subalpine conifers (Mayr et al., 2006). The gradual decrease in PLC was linked to the refilling of embolized tracheids (Figs. 4 and 5). Closed pits were rarely found during this period, indicating that pit aspiration recovered more rapidly than xylem embolism. Xylem refilling was suggested to be related to the absorption of water derived from rainfall and/or melting snow on the epidermal surfaces of needles and stems (Laur & Hacke, 2014; Losso et al., 2021; Mayr et al., 2014). Water may move via xylem and/or phloem from needles to stem xylem or via ray parenchyma from the stem surface to its inner tissues. As shown in stems sampled in late April 2017 (Fig. 4E), water-filled tracheids in earlywood were found only in tracheids near leaf traces, indicating that refilling occurred likely by water taken up over needles. The proposed refilling process may be based on osmotically induced positive pressure, which enables the resolving of air in water films within the tracheid lumens (Nardini et al., 2011; Ooeda et al., 2017) and finally also facilitate pit deaspiration (Mayr *et al*., 2014). Previous studies reported that PLC in conifer trees that did not incur severe frost drought gradually increased in winter but showed a comparably rapid reduction in spring (Sperry et al., 1994; Sparks et al., 2001; Mayr et al., 2003; McColloh et al., 2011; Ogasa et al., 2019; Maruta et al., 2020, 2022). Fast PLC changes may be based on pit deaspiration upon the recovery of water potentials in xylem sap.

### Pit aspiration by sap freezing (laboratory observations)

Pit aspiration induced by sap freezing was also confirmed under experimental conditions (Fig. 7). Ice formation within the xylem is a dynamically and spatially heterogeneous process (Neuner et al., 2010; Charrier et al., 2017), which can directly move pit membranes or indirectly cause pit aspiration via generated water flows (see Introduction). With respect to the latter, sinks for water drainage, which are either embolized tracheids or dehydrated parenchyma cells, will thereby allow sufficient flows for inducing pit aspiration in drought-stressed samples at low water potential. In contrast, at less negative xylem water potentials, the small (or absent) water sinks will limit the generation of pressure differences and the respective water flow over pits (Fig. 6). This is because, following Pascal’s principle, any pressure will first be transmitted equally to water in all xylem tracheids without water movement. Robson and Petty (1987) measured the pressures in the xylem sap during stem freezing and detected higher positive pressures in rehydrated than dehydrated stem xylem. This suggested that numerous spaces for water sinks (and thus for water flows) in dehydrated stems enabled a relaxation of pressures. Water movement to embolized vessels was observed in the midvein of snow gum (*Eucalyptus pauciflora*) while the leaf was freezing (Ball et al., 2006). Accordingly, reduced conductivity after freeze-thaw cycles was observed in dehydrated samples, whereas saturated samples were not affected (Fig. 6; Sperry et al., 1994; Feild & Brodribb, 2001; Pittermann & Sperry 2003; Mayr et al., 2003; Willson & Jackson, 2006; Ogasa et al., 2019; Maruta et al., 2020). Additionally, soaking in liquid N_2_ makes the stem segment rapidly but not completely homogeneously frozen. Closed and partly open pits found in dehydrated stem segments before freeze-thaw cycles might indicate artificial pit movements by cryofixation (Fig. 7A).

The water injection experiment demonstrated that loss of stem conductivity due to pit aspiration did not rapidly recover after the applied pressure was released (Fig. 8; Fig. S2; Sperry & Tyree, 1990). This result implied that the aspirated state of the pit-membrane was retained in the water-filled xylem while the experimental time was short (∼10 min). Also, in *A. veitchii* stems sampled from the field, the aspirated state of the pit-membrane was maintained at least until stem conductivity measurements after thawing. This suggested that closed pit membranes adhered to the cell wall of pit borders, in contrast to Hammel’s hypothesis, which expected pit membranes to elastically return to the central position directly after thawing (Hammel, 1967). In pits aspirated upon sap freezing, the torus may be strongly pressed to the cell walls of pit borders preventing rapid deaspiration. One study has suggested that pit-membrane is attached to the secondary wall of pit border by hydrogen bonding between polysaccharides and/or lignin hydroxyl groups in dried wood of *Pinus taeda* (Thomas & Kringstad, 1971). Because of larger contact area, a warty layer well developed on the surface of *Abies* tracheid cell walls (Fig. 1C) may be responsible for a tight adhesion in the overlap region of the torus and the pit border (Kohonen, 2006). However, the pit membranes returned from an aspirated position to a central position during overnight soaking of stems under partial vacuum (Fig. 7), indicating that deaspiration occurs within some hours in a pressurized situation. While xylem was refilled by vacuum infiltration, slow water flow likely occurred between tracheids, which detached the pit-membrane from the pit borders, so that the elasticity of margo microfibrils moved the torus to the central position. The pit-membrane flexibility of *A. veitchii* was estimated by the pressure gradient, inducing a 50% decrease in stem conductivity (3.65 kPa/mm; Fig. 8). It was similar to *P. abies* and *Pinus sylvestris* (4.66 and 3.35 kPa/mm, respectively) but much lower than *Juniperus communis* (13.1 kPa/mm; Beikircher et al., 2010). Pit membranes cannot move to an aspirated position unless the pressure generated during freezing of xylem sap is lower than the mechanical threshold (Hacke et al., 2004). Thus, there might be some species specificity on how fast pits aspirate (and deaspirate), and the effect of the mechanical flexibility of the margo should be examined on blockage of xylem water transport due to freezing-induced pit aspiration in future study.

## Conclusions

Winter xylem dysfunction in *A. veitchii* trees resulted from the combination of pit aspiration and xylem embolism. Whereas pit aspiration caused by freezing was responsible for xylem dysfunction during early winter when xylem water potentials were still moderate, embolism was the main factor limiting xylem water transport in mid and late winter. During recovery from xylem dysfunction, pit deaspiration occurred more rapidly than refilling of embolized tracheids. Overall, this combination led to an extended period of xylem dysfunction, which is likely relevant to many evergreen conifers exposed to winter drought and frequent freeze-thaw events. Future studies are required to better understand the interrelations among freeze-thaw- and drought-induced pit aspiration and embolism and the processes during embolism repair and pit deaspiration. Detailed insights into pit architecture, functioning, and the respective changes during ice formation in the xylem are prerequisites.

## Materials and Methods

### Plant materials and study sites

This study was performed on seven *A. veitchii* trees growing at a wind-exposed site near the Amaike Pass in the Yatsugatake ranges, central Japan (Chino City, Nagano Prefecture; 36°04′45″N, 138°19′57″E, 2250 m above sea level; Fig. 2A). Previous studies have reported that young stems lost hydraulic conductivity in winter and regained it until summer in *A. veitchii* growing at this site (Ogasa et al., 2019). The trees were ∼5 m high and exhibited a flag-shaped canopy that developed more leafy branches in the leeward relative to the windward direction (Fig. 2BC). Measurements were conducted between September 2016 and July 2017. Air temperatures were monitored with a datalogger (TR-52i; T&D Corp., Matsumoto, Japan). The number of freeze-thaw events was estimated from air temperature, assuming that freezing of xylem sap occurred between −1°C and −2°C (Fig. S1).

### Shoot water potential and stem hydraulic conductivity

For the seasonal course of shoot water potential and hydraulic conductivity, 5-year-old branches (Fig. 2c) were collected from five to seven wind-exposed trees (sampling dates: September 20, 2016; February 7, 2017; March 1, 2017; March 25, 2017; April 28, 2017; May 31, 2017; June 29, 2017; and July 27, 2017). One branch per tree was sampled around predawn, except for February to March, when the branches were at subzero temperatures even during daytime and expected to show little diurnal changes in water potential. All collected branches were put in a plastic bag, and shoot water potential was measured after transport to the laboratory on the same day. The remaining branches were stored at 4°C, and the hydraulic conductivity of the stems was measured on the day after sampling. In February and March, the collected and freezing branches were sent to the laboratory via refrigerated shipping (less than −20°C) to avoid thawing during transport. On the day after sampling, the collected branches were completely thawed at 4°C, and the shoot water potential and hydraulic conductivity of the stems were measured. Water potential was determined with a pressure chamber (Model 3000; Soilmoisture Equipment Corp., CA, USA). The measurement was made on current-year shoots of lateral branches bearing from a 2-year-old node.

For hydraulic measurements, the 1-year-old internode of the main branch was used. Needles were removed from the stem, and a section (∼3 cm long) was cut underwater to relax the xylem tension. From the distal part of the remaining internode, another 1-year-old 3-cm segment was cut underwater, soaked in liquid N_2_ for cryofixation, and stored at −80°C until observation with a cryo-SEM. Hydraulic conductivity (i.e., the ratio of flow rate to pressure gradient through the segment; *k*_h_) was measured by gravity-induced flow (Sperry et al., 1988). The 3-cm-long stem segment was girdled 5 mm from both cut ends to block the potential flow over the phloem. Both cut ends were trimmed with a razor blade underwater and attached to a tubing system. Water flow through the segment was generated by ∼5 kPa positive hydrostatic pressure from a reservoir of deionized water, and the flow rate was quantified gravimetrically with an electronic balance (AUW220; Shimadzu, Kyoto, Japan; measurements every 10 s for 1 min). To obtain maximum conductivity, the segments were submerged in deionized water (MilliQ Direct, Merck, Darmstadt, Germany) and exposed to partial vacuum (about −0.080 MPa) overnight (McCulloh et al., 2011). This procedure caused the refilling of embolized tracheids, and the flow rate was remeasured. To evaluate the inhibition of stem water transport, PLC (%) was calculated as

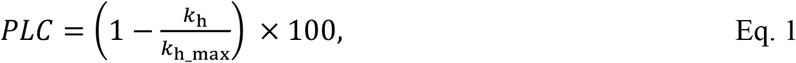

where *k*_h_ and *k*_h_max_ are the hydraulic conductivity before vacuum refilling and the maximum conductivity, respectively.

### Vulnerability curves

The vulnerability of *A. veitchii* stems to freeze-thaw-induced xylem dysfunction was estimated from plots of the percent loss of conductive area in xylem (PLC_area_) versus xylem water potential (referred to as vulnerability curve). Stems were subjected to 15 freeze-thaw cycles, and the resulting stem conductivity losses were quantified by the dye perfusion method (Hietz et al., 2008); PLC_area_ was calculated as an estimate for conductivity losses. Four 5-year-old branches were collected in November 2015 and dried to different water potentials between −0.5 and −2.5 MPa at ∼20°C on the benchtop. The branches were wrapped in a plastic bag and exposed to 15 freeze-thaw cycles between −9°C and 6°C [rate of temperature change: 3.6 ± 1.2°C/h, mean ± standard deviation (SD); duration of one cycle: 8 h] in a programmed incubator (FMU-133; Fukushima Industries Corp., Osaka, Japan). Stem freezing was observed between −1°C and −2°C, as indicated by exotherms. The 1-year-internode was cut to 3-cm-long segments underwater. The segment was girdled 5 mm from both cut ends and trimmed at both ends soaking in water. The basal end of the segment was connected to a tubing system filled with 0.1% acid fuchsine solution (Fujifilm Wako Pure Chemical Corp., Osaka, Japan) and filtered with a 0.22 μm membrane filter. The distal end was attached to a silicone tubing set under partial vacuum (about -5 kPa), and the fuchsine solution was perfused from the basal end for 10 min. The stained segment was immediately put in liquid N_2_ and dried with a freeze-dryer (FDU-2100; Eyela, Tokyo, Japan) for 2 days. After the dried segment was cut at 2 cm from the basal end, the cut surface was trimmed with a razor blade and digitalized with a digital camera (Moticam 2300; Shimadzu, Kyoto, Japan) mounted on a stereomicroscope (SZX16; Olympus, Tokyo, Japan). The stained and entire parts of the current-year annual ring were outlined using image editing software (GIMP 2.6; The GIMP Development Team). The xylem areas were measured using image analysis software (ImageJ; National Institutes of Health; http://imagej.net/contribute/citing). PLC_area_ was calculated by

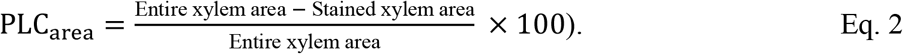

The relationship between PLC_area_ and xylem water potential was fitted via a sigmoidal function (Pammenter & Vander Willingen, 1998):

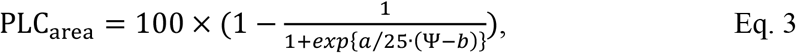

where Ψ is the xylem water potential, constant *a* is the slope at the inflection point, and *b* is xylem water potential that caused 50% loss of conductive area (*P*_50_area_). To assess the effect of freeze-thaw cycles on vulnerability to xylem dysfunction, the measured vulnerability curve for 15 freeze-thaw cycles was compared to vulnerability curves of stems subjected to drought or drought followed by 10 freeze-thaw cycles, as obtained previously by Ogasa *et al*. (2019).

### Observation of embolism and pit-membrane positions in stem xylem

The distribution of embolized tracheids and pit-membrane positions were observed with a cryo-SEM following the method of Yazaki et al. (2019). Frozen stem segments were carefully cut at ∼1 cm from the basal end. The new transverse cut surface was trimmed at ∼1 mm thickness with a cryostat (CryoStar NX70; Thermo Fischer Scientific K.K., Tokyo, Japan). The trimmed segment was fixed on a metal stage with an OCT compound (Sakura Finetek Japan, Tokyo, Japan) and positioned in a cryo-SEM chamber (JSM-6510 attached cryo-SEM unit; JEOL Ltd., Tokyo, Japan). Cryo-SEM observations were conducted at −130°C stage temperature at 1 to 3 kV accelerated voltage. The number of embolized tracheids and the positions of pit-membrane in pits were counted in four parts of the outermost annual ring formed in 2016 in the 1-year-old xylem (except for the un-mature annual ring formed in 2017): adaxial (normal wood), abaxial (reaction wood), and right and left sides. The percentage of embolized tracheids in 210 to 300 conduits located between rays was calculated for each part (Fig. 1A). Pit-membrane positions were observed in pits between water-filled tracheids. In 52 to 280 pits per sample, pit-membrane positions were clearly detectable and classified into three states: open, partly open, and closed (Fig. 1D). The “open” state was defined as a pit with the torus located in the central position within the pit chamber and the margo not attached to the pit borders. “Closed” pits exhibited the torus and margo completely attached to the pit border, which sealed the entire pit aperture. “Partly open” pits showed an intermediate position of the pit-membrane between open and closed states.

### Effect of freeze-thaw events on pit aspiration

To examine whether experimental freeze-thaw cycles cause pit aspiration in stem xylem, six 5-year-old branches were sampled from *A. veitchii* trees growing near the Amaike Pass on April 23, 2019, put in a plastic bag, and transported to the laboratory. The collected branches were wrapped in a plastic bag, and their cut ends were put in water at 4°C for 3 days for rehydration. The branches were dried at ∼20°C at the benchtop until the water potential of a current-year shoot reached −2 MPa (−2.0 ± 0.087 MPa). The water potential was measured by the pressure chamber method after equilibration of xylem pressure by wrapping the branches in a plastic bag for 0.5 to 1 h. To assess the pit-membrane position before experimental freeze-thaw treatment, a 3-cm-long stem segment of the 1-year-old internode was cut from the dried branches and used for cryo-SEM observations. In the following, the remaining dried branches were exposed to 11 freeze-thaw cycles from −6°C to 6°C in the programmed incubator (rate of temperature change: 1.68 ± 1.45°C/h, mean ± SD; duration of one cycle: 12 h). Exotherms indicated freezing between −1°C and −2°C (Fig. S1). From the 1-year-old internode of the treated branches, the stem segment was cut to 3 cm in length, and PLC was measured by gravity-induced flow, as described above. The remaining part of the internode was used for cryo-SEM observations to obtain the fraction of closed pits in the freeze-thaw-treated stems. Further, cryo-SEM observations were also conducted on the vacuum-infiltrated stem segments used for PLC measurements to analyze the repositioning of pit membranes to the open state.

### Pit aspiration induced by water injection

Pit aspiration induced by flow at high positive pressure was examined by measuring the flow rate over a sample under increasing and decreasing hydrostatic pressure. Rapid water flow on a conifer stem reduced the hydraulic conductivity, which is a ratio of flow rate to pressure gradient, probably due to pit closure (Beikircher et al., 2010; David-Schwartz et al., 2016; Sperry & Tyree, 1990). Six branches were collected from *A. veitchii* trees growing near the Amaike Pass on November 20, 2018. A 4-cm-long segment was cut from the 1-year-old internode. The cut end of the segment was trimmed underwater with a razor blade and connected to a high-pressure flow meter (Yang & Tyree 1994; Taneda et al., 2016). Water flow through the segment was generated at different positive pressures, and changes in hydraulic conductivity were monitored. The pressure gradient (=applied pressure/segment length) was recorded for 5% and 50% decreases in stem conductivity as a measure of the mechanical flexibility of pit membranes. When the stem conductivity decreased to <30% of its initial value, the applied pressure was gradually released and the stem conductivity was measured at the relaxed pressure.

### Statistical analyses

Seasonal changes in xylem water potential, PLC, fractions of pit-membrane positions, and embolized tracheids were tested by analysis of variance (ANOVA), followed by a post-hoc test for pairwise differences using Tukey’s corrections for multiple comparisons and a significance threshold at *P* < 0.05. The significance of the difference between fractions of three pit-membrane states and between embolized tracheids in different parts of xylem were also tested for each month by ANOVA, followed by a post-hoc test for pairwise differences at *P* < 0.05 (Tables S1 and S2). Likewise, to assess the effects of experimental freeze-thaw cycles and rehydration under a partial vacuum, the significance of differences in the fractions of pit-membrane states was tested by ANOVA and a post-hoc test for pairwise differences at *P* < 0.05. A vulnerability curve for *A. veitchii* stems subjected to 15 cycles of freeze-thaw events was regressed to a logistic function (Eq. 2) by the least-squares method. The relationship between PLC and a fraction of closed pits from the experimental freeze-thaw treatment was tested by Pearson’s correlation test. Through the text, when averages of the measured values are mentioned, the shown values are the mean ± 1 standard error (SE). All statistical analyses were performed using R version 4.0.3 (R Core Team, 2020).

## Acknowledgments and Funding

We thank Drs. I. Terashima, Y. Sano, K. Kuroda, and K. Fukuda for invaluable discussion. This study was financially supported by JSPS KAKENHI (grant no. 17H03825) and Moriya research grant. This study was also supported by the Austrian Research Agency (FWF) project P32203 and I4918-B and the research area “Mountain Regions” of the University of Innsbruck.

## Author Contributions

HT, MO, KY, and EM planned and designed the research. HT, MO, KY, and YM conducted fieldwork and measurements in the laboratory. HT and SNF conducted image analyses. HT, SM, MO, KY, YM, and EM contributed to the interpretation of the data and wrote the manuscript.

## Supplemental Data

**Figure S1.**
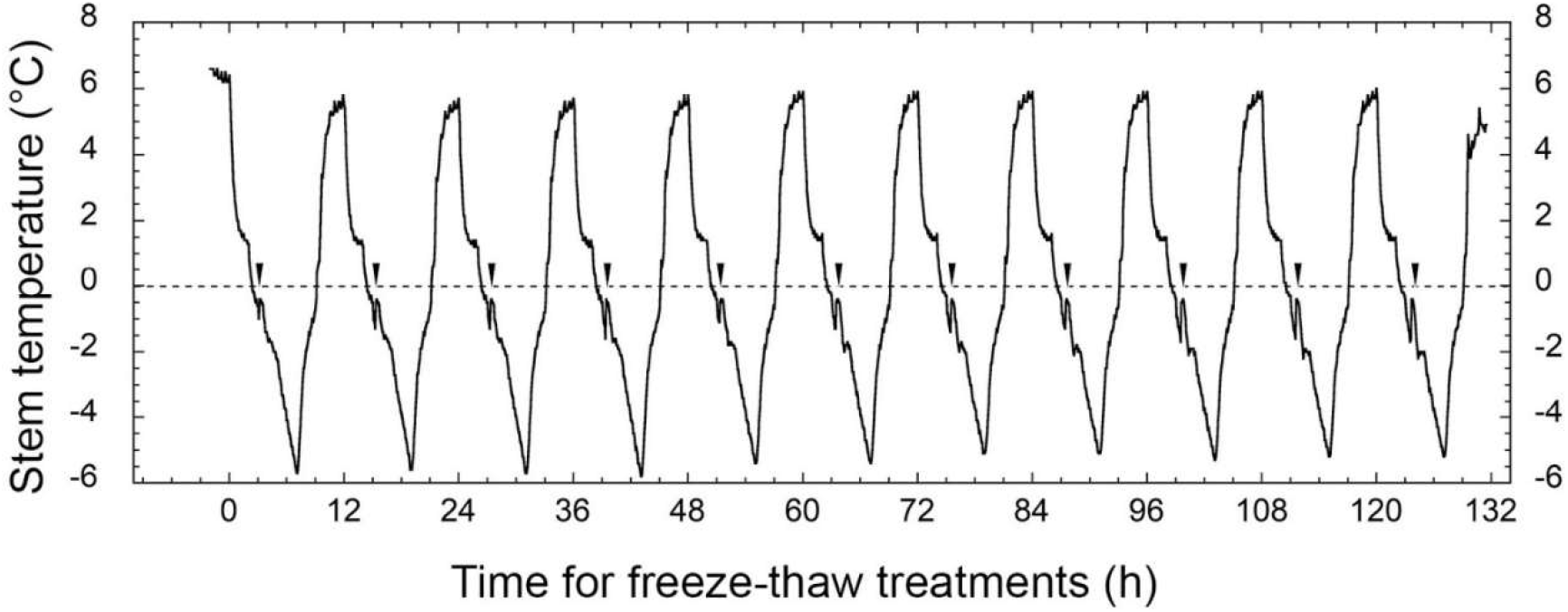
Change in stem temperature during experimental freeze-thaw cycles. A probe of thermo-couples was put in the bark of the 1-year-old internode. Arrowheads represent the occurrence of exotherms.

**Figure S2.**
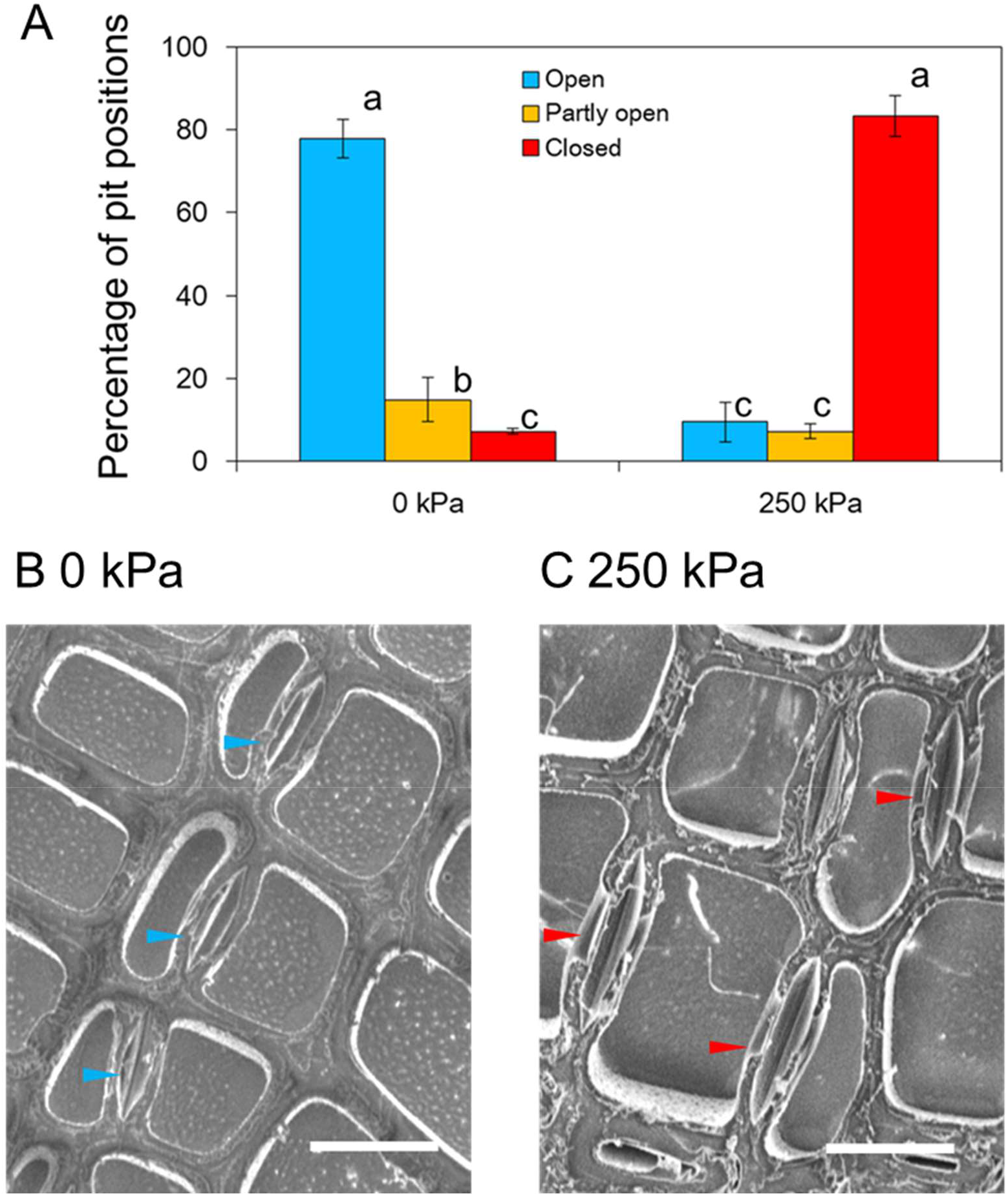
Changes in the pit-membrane position via water flow at different pressures in *A. sachaliensis*. (A) Percentage of the three pit-membrane positions: open, partly open, and closed. Pit membranes in xylem (B) before and (C) after an injection of water flow pressurized at ∼250 kPa. Six stem segments of 1-year-old internodes of *A. sachaliensis* were collected in the early morning from a potted plant grown at a sunny field at the University of Tokyo, cut to ∼4 cm, and soaked in water under −0.08 MPa of a partial vacuum overnight to refill xylem with water and restore pit membranes to the central positions. Half of the six stem segments were frozen with liquid N_2_, and the remaining stem segments were attached to a high-pressure flow meter, and water was injected as the applied pressure periodically increased at an interval of ∼30 to 60 kPa to ∼250 kPa. When water flow was injected to the stem segments at ∼250 kPa (∼6.25 kPa/mm; the stem conductivity reduced by ∼80% of that at 30 kPa), the segment was cryofixed using liquid N_2_. After trimming the cut end with a cryostat, the frozen stem segments were observed with a cryo-SEM at 3 kV of accelerated voltage and about −120°C of stage temperature. A total of 120 and 121 pits located in normal wood from three stems were counted for control (0 kPa) and pressurized stems (250 kPa), respectively. Error bars in (A) represent ±1 SE (*n* = 3). Letters beside the columns in (A) represent significant differences by ANOVA and Tukey’s multiple comparison tests at *P* < 0.05. Blue and red arrowheads in (B and C) represent open and closed pit membranes, respectively. Scale bar, 10 μm (B and C).

**Table S1.**
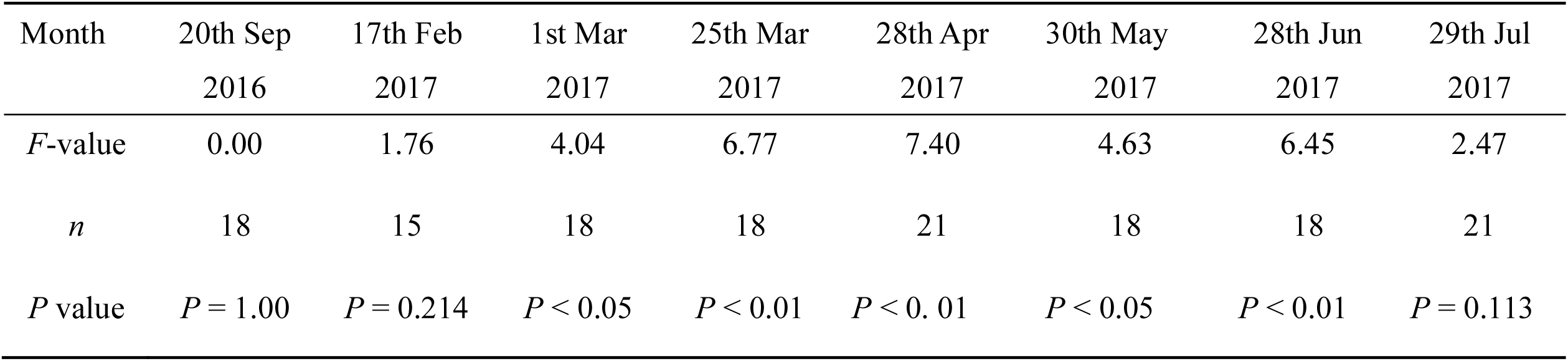
The significant difference in a fraction of embolized tracheids among normal and reaction wood in *Abies veitchii* trees and the mean by ANOVA and significant threshold at *P* < 0.05.

**Table S2.** The significant difference in a fraction of three pit positions (open, partly open, closed) in *Abies veitchii* trees and the mean by ANOVA and significant threshold at *P* < 0.05.

| Month | 20th Sep<br>2016 | 17th Feb<br>2017 | 1st Mar<br>2017 | 25th Mar<br>2017 | 28th Apr<br>2017 | 30th May<br>2017 | 28th Jun<br>2017 | 29th Jul<br>2017 |
| --- | --- | --- | --- | --- | --- | --- | --- | --- |
| *F*-value | 330 | 334 | 391 | 5.68 | 24.2 | 61.1 | 33.7 | 103 |
| *n* | 12 | 15 | 18 | 18 | 21 | 18 | 18 | 12 |
| *P* value | $P < 0.001$ | $P < 0.001$ | $P < 0.001$ | $P < 0.05$ | $P < 0.001$ | $P < 0.001$ | $P < 0.001$ | $P < 0.001$ |

